# An enhanced molecular tension sensor based on bioluminescence resonance energy transfer (BRET)

**DOI:** 10.1101/617696

**Authors:** Eric J. Aird, Kassidy J. Tompkins, Wendy R. Gordon

## Abstract

Molecular tension sensors measure piconewton forces experienced by individual proteins in the context of the cellular microenvironment. Current genetically-encoded tension sensors use FRET to report on extension of an elastic peptide encoded in a cellular protein of interest. Here we present the development and characterization of a new type of molecular tension sensor based on bioluminescence resonance energy transfer (BRET) which exhibits more desirable spectral properties and an enhanced dynamic range compared to other molecular tension sensors. Moreover, it avoids many disadvantages of FRET measurements in cells, including heating of the sample, autofluorescence, photobleaching, and corrections of direct acceptor excitation. We benchmark the sensor by inserting it into the canonical mechanosensing focal adhesion protein vinculin, observing highly resolved gradients of tensional changes across focal adhesions. We anticipate that the BRET-TS will expand the toolkit available to study mechanotransduction at a molecular level and allow potential extension to an in vivo context.

## INTRODUCTION

The mechanical microenvironment of a cell guides cellular processes such as migration, differentiation, division, and signaling^1^, and is thus often altered in disease^2,3^. At a molecular level, mechanical stimuli alter conformations of mechanosensing proteins to communicate the stimulus to the cell interior and effect a downstream cellular response in a process called mechanotransduction^4,5^. Measuring tensions sensed by cellular proteins and how tensions are altered by cellular context will lead to better understanding of the molecular mechanisms used in mechanosensing in disease-related processes such as tumor migration.

Tools to measure tensions at a cell and tissue level such as traction force microscopy and atomic force microscopy are well-established. Tools to measure molecular level tensions have only emerged more recently to characterize the pN forces present across mechanosensitive proteins^6^ and have been used to measure tensions sensed by focal adhesions proteins such as vinculin, talin, and integrins, as well as cadherins^7–10^. Generally, genetically encodable molecular tension sensors contain an elastic peptide linker with well-characterized extension in response to applied force flanked by two fluorescent proteins (FP) allowing for measurement of tension across a protein typically utilizing Förster resonance energy transfer (FRET). The extent of linker stretching due to mechanical force can measured by FRET and subsequently be correlated to a quantifiable force. Until recently, the state of the art genetically-encoded FRET tension was TSMod, a distance dependent FRET-based sensor that can measure tensions in the 1-6 pN range^7^. The dynamic range of TSMod, however, is limited by the photophysical properties of the fluorescent protein pair, mTFP1 and venus(A206K)^11^. Recent studies reported improved FRET tension sensors by using more optimal fluorescent proteins, utilizing different linkers, and engineering FP termini to achieve closer distances that yielded higher FRET under no load^12,13^. A computational model has also been described to allow researchers to optimize force range or sensitivity based on experimental needs^12^. Nonetheless, these sensors are still constrained by the limitations of FRET, namely autofluorescence, direct acceptor excitation, photobleaching, and incompatibility for *in vivo* use.

We developed a genetically-encodable molecular tension sensor based on bioluminescence resonance energy transfer (BRET) that overcomes several limitations of FRET. BRET is a distance and orientation dependent phenomenon analogous to FRET but is initiated by a chemiluminescent reaction of a luciferase protein with its substrate instead of light excitation^14^. Upon luciferase excitation, non-radiative energy transfer can occur to excite a closely linked acceptor fluorescent protein that produces a distinct emission spectrum. Our BRET tension sensor, which we will refer to as BRET-TS, takes advantage of the uncharacteristically bright Nanoluciferase (NanoLuc), allowing in vitro and live cell detection of signal using standard plate readers and microscopes. The luminescent-fluorescent protein pair of NanoLuc^15^ and mNeonGreen^16^ has recently been shown to exhibit robust resonance energy transfer efficiency^17^. Here we report on using these proteins as a sensitive BRET donor-acceptor pair in a molecular tension sensor. Using BRET as a distance reporter in molecular tension sensors allows for more sensitive readouts due to the lack of autofluorescence and phototoxicity. BRET also offers the potential for use in vivo where light excitation is not able to penetrate into tissue. We demonstrate the utility of this tension sensor in vitro and in cell based assays, using vinculin as the model system to show an enhanced dynamic range compared to field standard FRET-based sensors.

## RESULTS AND DISCUSSION

### Sensor design and characterization

In the BRET-based molecular tension sensor (BRET-TS), the NanoLuc donor and mNeonGreen acceptor proteins flank a 40 amino acid flexible spider silk flagelliform domain (GPGGA)_8_ (Figure 1A). Spider silk exhibits predictable changes in length under tension^18^. Both proteins are terminally truncated to enhance the base BRET efficiency by bringing them in closer proximity^17^. We first expressed recombinant BRET-TS and the commonly used FRET-TSMod to compare their spectral properties^7^.

**Figure 1.**
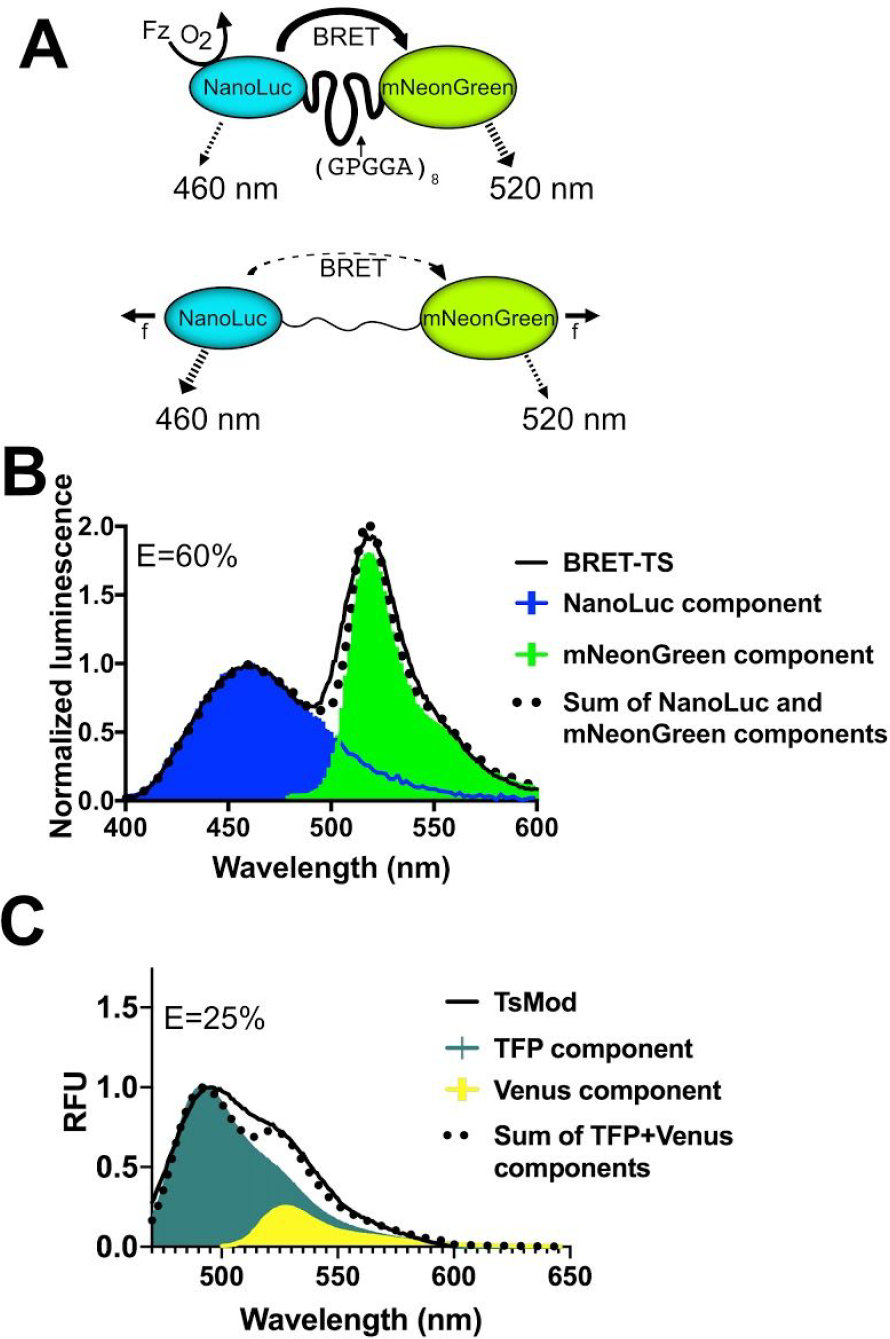
Schematic of the genetically-encodable BRET molecular tension sensor (BRET-TS). (A) NanoLuc, the energy donor, is excited via addition of furimazine (Fz) in the presence of oxygen. In the absence of force (f), resonance energy transfer occurs to the fluorescent acceptor, mNeonGreen. With applied force, the donor-acceptor pair is separated, reducing resonance energy transfer. Unloaded spectral and resonance energy transfer properties of (B) BRET-TS in comparison to (C) FRET-based TSMod. Spectral components and resulting best-fit additive spectrum depicted. Axes in units of luminescence normalized at 460nm and relative fluorescence units (RFU). Energy transfer efficiencies E were calculated as described in Methods.

In the absence of tension, the unloaded BRET sensor boasts a robust ~60% BRET efficiency upon addition of NanoLuc’s chemiluminescent substrate, furimazine (Figure 1B). Moreover, the emission maxima of NanoLuc (460 nm) and mNeonGreen (517 nm) are spectrally separated by 57 nm. In contrast, the unloaded TSMod FRET tension sensor exhibits only 25% energy transfer with extensive spectral overlap between mTFP and Venus (Figure 1C)^7^. Recent iterations of FRET tension sensors using Clover and Ruby derived FPs display improved spectral separation but still only achieve unloaded energy transfer efficiencies of 35%.^12^

A BRET efficiency of 60% approaches energy transfer efficiencies observed for unloaded spider silk peptides flanked by organic fluorophores Cy3 and Cy5^7,26^ used to calibrate the sensors, despite the usually detrimental contribution of increased distance that FPs contribute. This is likely due to the fact that in BRET-TS, the mNeonGreen acceptor has a higher quantum efficiency than the NanoLuc donor^17^, a trend that is reversed in the mTFP1 / venus and clover / ruby FRET pairs. The enhanced BRET efficiency at zero force translates to a larger dynamic range, as the sensor has the potential to detect a 60% change in energy transfer when force is applied, making it easier to detect higher forces and resolve the entire range of forces. The large spectral separation of the donor and acceptor is also desirable for ratiometric imaging, where spectral filters are used to separate emission from donor and acceptor.

Upon application of tension to the elastic peptide, we expect the spider silk linker to stretch, separating the protein pair and causing a decrease in the resonance energy transfer (RET) efficiency. Thus, we next aimed to ensure that BRET-TS exhibits the expected changes in RET with distance between the donor and acceptor. We initially characterized the distance dependence of BRET by inserting a series of rigid alpha helical linkers (HL) A(EAAAK)_n_A (Figure 2A) of known lengths between mNeonGreen and NanoLuc in the context of recombinant proteins expressed in *E. Coli*^19^. Expectedly, as the distance between the protein pair increases, a decrease in BRET is observed when measured in bulk using a standard plate reader (Figure 2B).

**Figure 2.**
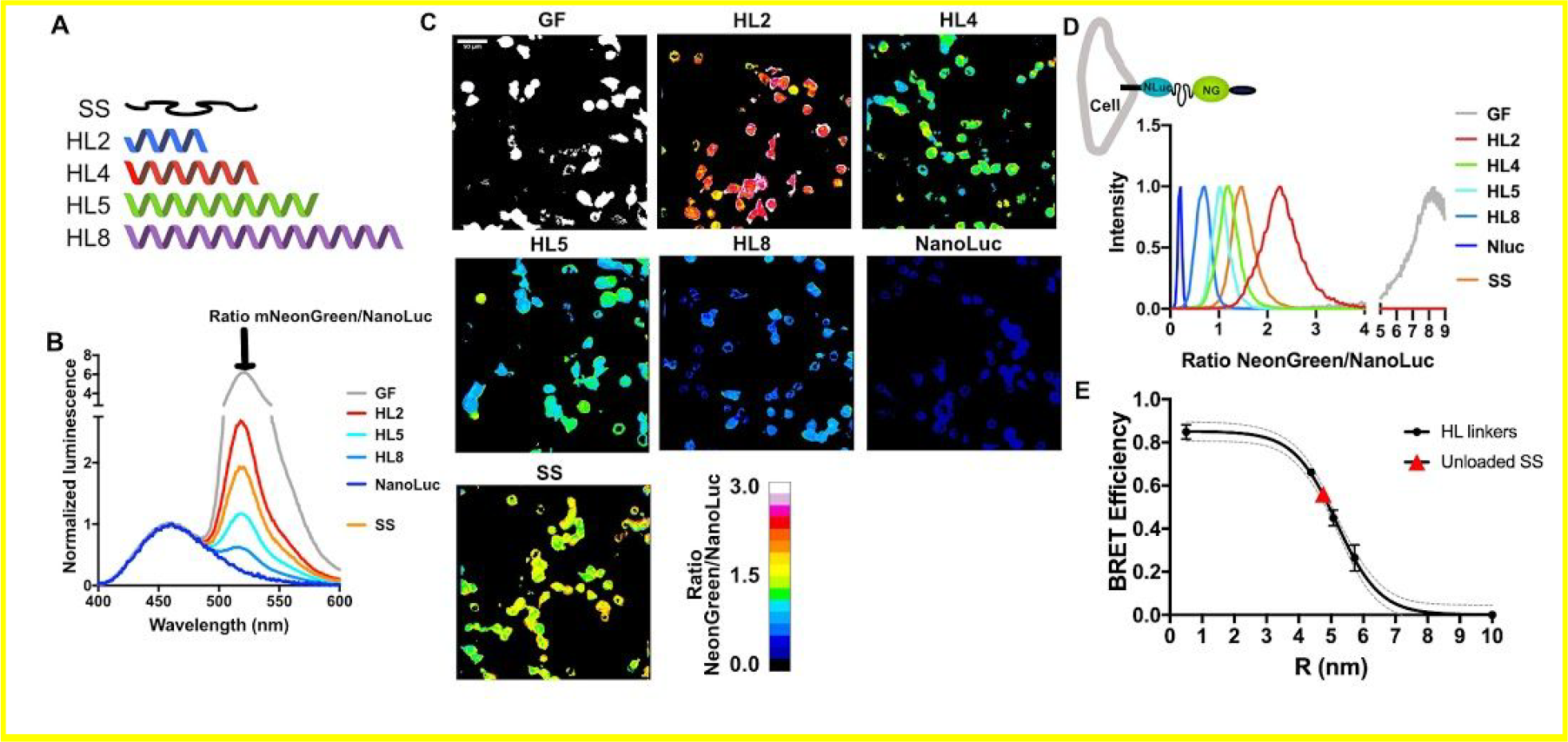
Characterization of distance dependence of BRET-TS in vitro and in cells. (A) Rigid alpha helical linkers (HL) of varying lengths were inserted between the mNeonGreen-NanoLuc pair^19^ in addition to a minimal dipeptide GF linker. NanoLuc alone was also measured. (B) Spectral plate reader emissions of recombinantly expressed proteins, normalized to NanoLuc peak emission (460 nm). (C) HEK293T cells transfected with the various HL linker constructs inserted into a plasma membrane localized protein. Cells were imaged using NanoLuc and mNeonGreen specific spectral filters and ratiometric mNeonGreen:NanoLuc images are shown. (D) Histograms of a field of cells from C were calculated and normalized to 1 to derive the average BRET ratio for a given linker. (E) Sigmoidal curves comparing HL linker BRET efficiency derived from in vitro recombinant protein BRET-TS and cell-surface BRET-TS versus distance r, calculated from the BRET data and using the measured R_0_.

We next encoded the series of alpha helical linkers sandwiched between mNeonGreen and NanoLuc into a cell surface expressed protein to characterize RET as a function of distance in cells. Constructs were transiently transfected into HEK293T or U2OS cells, and BRET read out on a plate reader on via ratiometric imaging. Remarkably, the cellular BRET was easily detectable on a plate reader, underscoring the brightness and low background of NanoLuc (Figure S1). To detect BRET localization in live cells, furimazine was added to the media and the ratio of emission of mNeonGreen to NanoLuc was measured simultaneously or sequentially using an EM-CCD camera outfitted with spectral filters specific to NanoLuc and mNeonGreen emissions (Figure 2C). The pixel-by-pixel ratio of the two images was then calculated. Emission from NanoLuc alone as well as emission from the donor and acceptor connected by a short dipeptide “GF” linker were also measured for reference. Again, a drastic decrease in BRET is observed with increasing linker length (Figure 2D). Heterogeneity in the BRET signal, likely reflecting a diversity in mechanical microenvironments sensed by the cell-surface BRET-TS, is also apparent in the cell experiments that is masked in the in vitro bulk experiments.

To ensure the BRET observed was due to intramolecular rather than intermolecular effects, we co-transfected cell surface receptors containing either NanoLuc or mNeonGreen alone. We did not observe any BRET signal, consistent with a lack of intermolecular BRET in this context (Figure S2A). Similar results were obtained when inserting a 3C proteolytic cleavage site in the recombinant protein version. Cleavage by HRV 3C protease abolished BRET resulting in only the expected NanoLuc emission (Figure S2B).

To determine if the ratios derived from in vitro and in cell linker experiments were similar, we plotted the BRET efficiency as a function of distance. We first calculated histograms of ratiometric images from Figure 2B to determine average NeonGreen to NanoLuc ratios. To convert BRET ratios to BRET efficiencies, we first corrected for the Nanoluc/NeonGreen emission overlap and calculated BRET efficiency using spectral deconvolution or the ratio of intensities (Supplemental Methods).

We next calculated the Förster radius (R_0_) of mNeonGreen-NanoLuc to be 51 Å based on the experimentally determined spectral overlap (J=2.84*10^15^ nm^4^ M^−1^ cm^−1^) and using NanoLuc’s previously reported quantum yield^20^. This is the highest reported Förster radius of any NanoLuc-FP pair^21^. Using this R_0_ and our experimental BRET efficiencies, we then calculated R values for the helical linkers using standard FRET equations and plotted them against BRET efficiency (Figure 2E). The curves fit well to a sigmoidal function. A similar curve is obtained using the R values previously calculated for these linkers^19^ (Figure S3). The BRET efficiencies measured in vitro and in cells were remarkably similar for a given linker, and we observed no significant difference between the image splitter and filter wheel methods of ratiometric imaging.

### Measuring tension across vinculin

Finally, we wanted to validate BRET-TS in a known mechanosensing protein. We chose the focal adhesion protein vinculin that has previously been shown to experience tensions on the order of 1 to 6 pN using molecular tension sensors^7^. This system provides a good opportunity to benchmark the dynamic range of BRET-TS against state-of-the-art FRET tension sensors. We inserted the BRET-TS into a previously described site within the protein after residue 883 (VinTS). We also created a force-insensitive control construct lacking the carboxyl terminal F-actin binding tail (VinTL) (Figure 3A). As this tension sensor is 84 amino acids smaller than TSMod, we did not repeat previously reported functional characterization of vinculin with the inserted sensor^7^.

**Figure 3.**
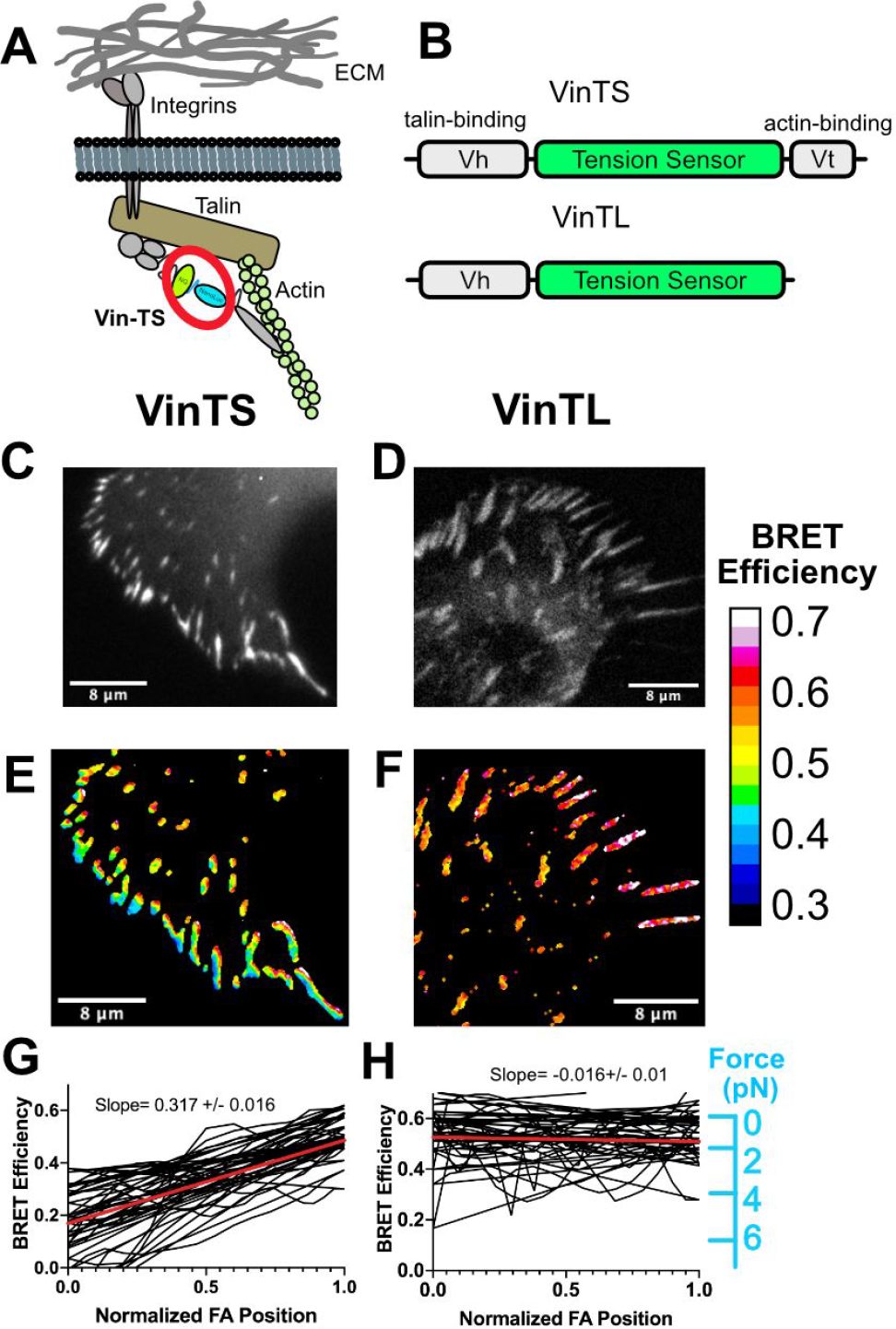
Measuring tension across vinculin. (A) Schematic of vinculin with inserted BRET tension sensor (BRET-TS) in the context of a focal adhesion. (B) Vinculin constructs used in these studies: full length vinculin with inserted BRET-TS (VinTS) and actin binding deficient vinculin with BRET-TS (VinTL). (C-D) Unprocessed luminescent images of focal adhesions. (E-F) Processed, ratiometric images from (C-D). Scale bar displayed as BRET efficiency (out of 1 maximum). (G-F) Manual line scans taken across focal adhesions, normalized for focal adhesion (FA) length (n=30). Images and line scans are representative from images collected over numerous independent experiments. Force is extrapolated from FRET vs force curves of spider-silk peptide labeled with Cy3/Cy5^26^

Remarkably, luciferase signal from VinTS was clearly observable in a bulk plate reader assay in comparison to zero signal in untransfected cells (Figure S4). In contrast, direct excitation of mNeonGreen at 450nm resulted in the same signal in both untransfected and transfected cells due to cellular autofluorescence. For imaging of VinTS in focal adhesions, constructs were transfected in U2OS cells and plated sparsely on fibronectin. The cells were subsequently imaged upon addition of furimazine. We observed a gradient in tension across peripheral focal adhesions^22^ in VinTS that was not observed in the force-insensitive mutant VinTL (Figure 3B-H). Tension gradients were not observed in the original FRET-TS TSMod due to insufficient dynamic range^7^, but were recently detected in improved FRET-TS’s^12^. BRET efficiency is lowest at the cell periphery, which corresponds to higher forces. BRET efficiency gradually increases (force decreases) moving towards the nucleus (Figure 3C). This gradient is abolished in the actin binding mutant lacking the carboxy-terminus of vinculin and in point mutants of the actin or talin binding domains (Figure 3H and S5)^23^. BRET-TS provides better resolution in highlighting the gradient across the focal adhesion compared to previous studies; we observe an average 31% change in BRET across peripheral focal adhesions, compared to 5% and 15% BRET gradients reported for TSMod and improved TSMods, respectively^7,12^. VinTS has already been used to dissect molecular mechanisms of collagen matrix formation in stem cells^24^ and measure the effect of osteocytes on migration potential of breast cancer cells^25^. The enhanced dynamic range of VinTS will allow further studies of focal adhesion tensions in cellular processes.

## CONCLUSIONS

We developed a genetically-encodable BRET-based tension sensor that is smaller in size, offers better spectral separation, and a larger dynamic range for detecting pN forces over FRET based molecular tension sensors due to the desirable photophysical properties of the mNeonGreen-NanoLuc pair^17^. Moreover, BRET offers other key improvements over FRET-based sensors such as lack of photo toxicity, greatly simplified image processing and increased signal-to-noise due to the use of bioluminescence as the energy donor. Though the enhanced dynamic range and potential for in vivo imaging of BRET-TS will be advantageous for most applications of molecular tension sensors, BRET-TS would not be suitable for measuring tensions related to cellular processes that happen in <10 seconds due to the longer exposure times required. Moreover, BRET-TS is not compatible with optical sectioning strategies utilizing laser scanning, such as confocal microscopy, and would necessitate the use of computational optical sectioning.

Further optimization of the force regime can be accomplished through substitution of the flexible spider silk flagelliform linker with other recently described molecular spring domains^13,26^. Furthermore, we believe this sensor will be aptly suited for live animal in vivo measurement of molecular scale tensions. Finally, this BRET pair should be widely applicable to use in other biosensors.

## Author Contributions

E.J.A., K.J.T. and W.R.G. designed experiments, collected data, and analyzed data. The manuscript was written through contributions of all authors. All authors have given approval to the final version of the manuscript.

## ACKNOWLEDGMENT

We would like to thank the Physical Sciences of Oncology Center at UMN for resources and support. Thanks to Fluorescence Innovations as well as Thomas Pengo of the University Imaging Centers for support and helpful discussions. Thanks to Brenton Hoffman for helpful discussions. We would also like to thank Hideki Aihara and Nick Levinson for use of their plate readers.

## ABBREVIATIONS

BRET: Bioluminescence resonance energy transfer
FRET: Förster resonance energy transfer
FP: fluorescent protein
TS: tension sensor
RET: resonance energy transfer
HL: helical linker
FA: focal adhesion
RLU: relative light units
Nluc: NanoLuc

## Methods

### Tension sensor construct design

The mNeonGreenΔC10-spider silk flagelliform-NanoLucΔN5 tension sensor and HL linkers DNA sequences were synthesized by Integrated DNA Technologies (IDT) as gBlocks containing overhangs to be cloned between HUH tags RepBm and mMobA into pTD68-SUMO-Protein G-RepBm-mMobA vector backbone via Infusion cloning (Takara Bio)^27^. The TSMod recombinant protein was made in the same context, except via PCR amplification of the TSMod DNA (Addgene #26019, courtesy of Martin Schwartz). For insertion into existing pcDNA3.1 mammalian expression vector containing a signal sequence, CD8 juxtamembrane sequence, and DLL4 transmembrane and intracellular domain, the tension sensor was PCR amplified and inserted using SbfI and BamHI restriction sites. Insertion into the vinculin locus (Addgene # 26019 and 26020, courtesy of Martin Schwartz) was performed using Golden Gate assembly. Vinculin point mutants I997A and A50I were constructed using QuikChange II Site-Directed Mutagenesis (Agilent).

### Protein expression and purification

Bacterial expression of proteins was performed in BL21(DE3) *E. coli* using an autoinduction protocol^28^. Following inoculation, cells were rotated at 300 rpm for 8 hours at 37°C followed by 24 hours at 24°C. Cells were pelleted and lysed in 50 mM Tris, 200 mM NaCl, 20% sucrose, pH 7.4. The clarified supernatant was first passed over Ni-NTA resin (Thermo Scientific), washed with 15 column volumes of 50 mM Tris, 200 mM NaCl, 15 mM imidazole, pH 7.4, and eluted with 250 mM imidazole followed by overnight incubation with SUMO protease Ulp1 at 4°C while dialyzing into 50 mM Tris, pH 7.4, 200 mM NaCl, 10 mM CaCl_2_, 1 mM DTT. Protein was concentrated in a 30 kDa MWCO spin concentrator (Amicon) and applied to a Bio-Rad 650 size exclusion chromatography column. Fractions were analyzed by SDS-PAGE and pooled. Concentration was measured by Bradford Assay (BioRad) or A_280_ absorbance. Concentrated aliquots were stored at −20°C or flash frozen in dry ice / IPA.

### Cell culture conditions

HEK293T and U2OS cells were cultured in DMEM (Corning) supplemented with 10% FBS (Gibco) and 0.5% penicillin / streptomycin (Gibco). Cells were incubated at 37°C in 5% CO_2_.

### Transfections

Transfections were carried out with Lipofectamine 3000 (Invitrogen) using the manufacturer’s suggested protocol. Cells were plated on untreated or poly-lysine glass bottom 96-well plates (MatTek). 24-48 hours post-transfection, cells were assayed by exchanging the media with phenol red-free DMEM (Corning) and adding 1:100 dilution of furimazine (Promega) to activate NanoLuc. For vinculin experiments, fibronectin (Sigma) was diluted in PBS and plated at 10 µg/ml 16 hours prior to transfection. Wells were washed twice with PBS before adding cells.

### 3C cleavage assay

Human Rhinovirus (HRV) 3C protease (5 units) was incubated with 3 µM purified protein containing a 3C cleavage site between mNeonGreen and NanoLuc in 50 mM Tris, pH 7.4, 200 mM NaCl, 10 mM CaCl_2_, 1 mM DTT for up to 24 hours. Reactions were analyzed on a plate reader at the given timepoints.

### BRET measurements

Both *in vitro* and cell-based plate reader measurements were performed in 96 well plates on either a Spark or M1000Pro (Tecan). On the M1000Pro, spectral scans were acquired in luminescence scanning mode. On the Spark, spectral scans needed to be acquired in fluorescence mode using the longest wavelength excitation light at 900 nm, well below the energy required to excite electronic transitions of mNeonGreen. Spectra were deconvoluted using Prism (GraphPad).

### Microscopy

Live cell imaging was carried out on an Olympus IX83 inverted microscope equipped with an Andor iXon Ultra 888 EM-CCD camera and a magnification changer to increase magnification of 60x images to 120x. Images were acquired in one of two methods, either using a W-view Gemini image splitter (Hamamatsu) with Semrock light filters FF01-460/60 (for NanoLuc) and Semrock FF01-536/40 (for mNeonGreen) or using a filter wheel with two separate acquisitions with Semrock light filters FF01-460/60 (for NanoLuc) and Semrock FF01-536/40 (for mNeonGreen). Integration times for both capture methods ranged from 15 seconds to 2 minutes.

### Image analysis

Images were analyzed on Fiji (version 1.51v). Stacked images were first aligned using the Linear Stack with Alignment with SIFT plugin. Images were then analyzed using the BRET-Analyzer plugin^29^. For vinculin experiments, images were analyzed using a custom Fiji macro. Briefly, images were masked for focal adhesions by averaging the intensity, rolling background subtraction, and thresholding. The mask was then applied to aligned images prior to division and smoothing. Intensity was visualized as 16 color LUT. Line scanning was carried out across individual focal adhesions by manually using the line tool, starting from the outer portion of the cell. Intensity profiles were then exported in text format, normalized to focal adhesion length, and graphed.

### Förster radius calculation

The emission spectrum of NanoLuc and the absorbance spectrum of mNeonGreen were experimentally obtained using a M1000Pro (Tecan). The spectral overlap component J was calculated (a | e 1.2, FluorTools) and entered into the Förster equation to determine the Förster radius (R_0_) of this truncated NanoLuc-mNeonGreen pair.

### Calculation of BRET efficiency from spectra and EM-CCD images

Spectra were fit as the sum of two Gaussians, with mean of the Gaussians constrained to 460 nm (NanoLuc emission) and 520 nm (mNeonGreen emission). BRET efficiency was calculated from the amplitudes using the equation: BRET efficiency= Amp_NG_/(Amp_NG_+Amp_NLuc_). These results were comparable to spectral deconvolution methods in which coefficients of mNeonGreen and NanoLuc spectra were varied to achieve best fit to a given emission spectrum. Finally, BRET efficiency was calculated a third way using corrected ratios of intensities: BRET efficiency= I_NG@520_-I_NLuc@520_/[I_NLuc@460_+(I_NG@520_-I_NLuc@520_)]. TSMod, the emission spectra of Venus and mTFP1 were spectrally too close to fit using two Gaussians. Unloaded FRET efficiency of 23% was calculated using spectral deconvolution using the equation FRET efficiency= Coeff_Venus_/(Coeff_Venus_+Coeff_mTFP_). For TsMod, the coefficients of mTFP and Venus were 1 and 0.3, resulting in a BRET efficiency of ~25%. For NanoLuc and mNeonGreen, the coefficients were 1 and 1.8 with a NLuc@520 of 0.3, resulting in a BRET efficiency of ~60%. Ratios of emission from mNeonGreen filter to NanoLuc filter for surface expressing tension sensors harboring different linkers were calculated using BRET efficiency= I_NG@NG_-I_NLuc@NG_/[I_NLuc@NLuc_+(I_NG@NG_-I_NLuc@NG_)]. For Vinculin images, the average ratio of VinTL was normalized to the average ratio of unloaded BRET-TS in the BRET vs R curve, and this normalization applied to VinTS data. After ratios were normalized, BRET efficiency was calculated using: BRET efficiency= I_NG@NG_/[I_NLuc@NLuc_+I_NG@NG_)]. Forces were extrapolated using previous calibrations of spider silk tension sensor flanked by Cy3 and Cy5, which had an almost identical unloaded resonance energy transfer efficiency.

**Figure S1.**
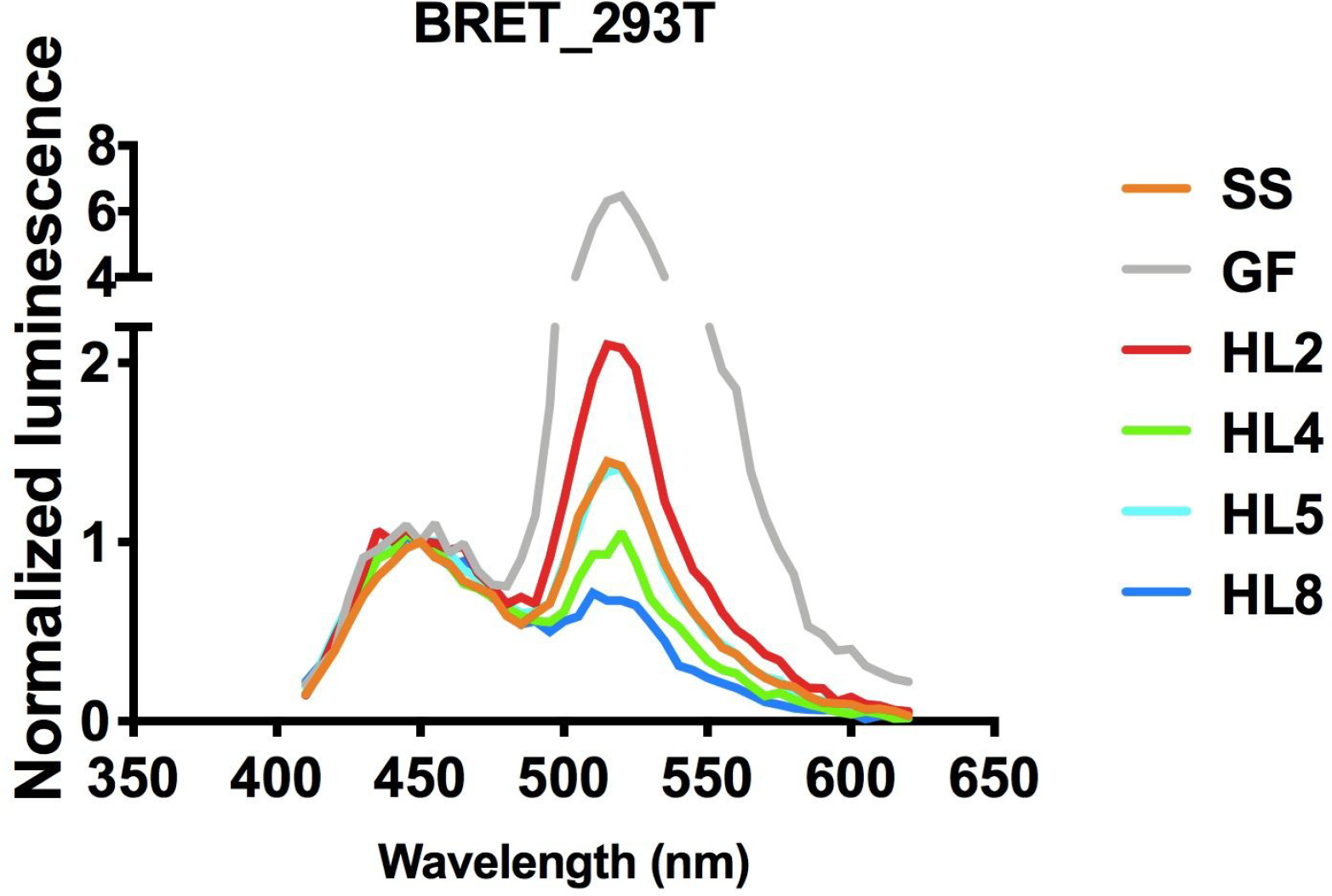
Readout of BRET of HL linkers inserted into cell-surface tension sensor and transfected in HEK293T cells in 96 well plates upon addition of Fz using spectral platereader

**Figure S2.**
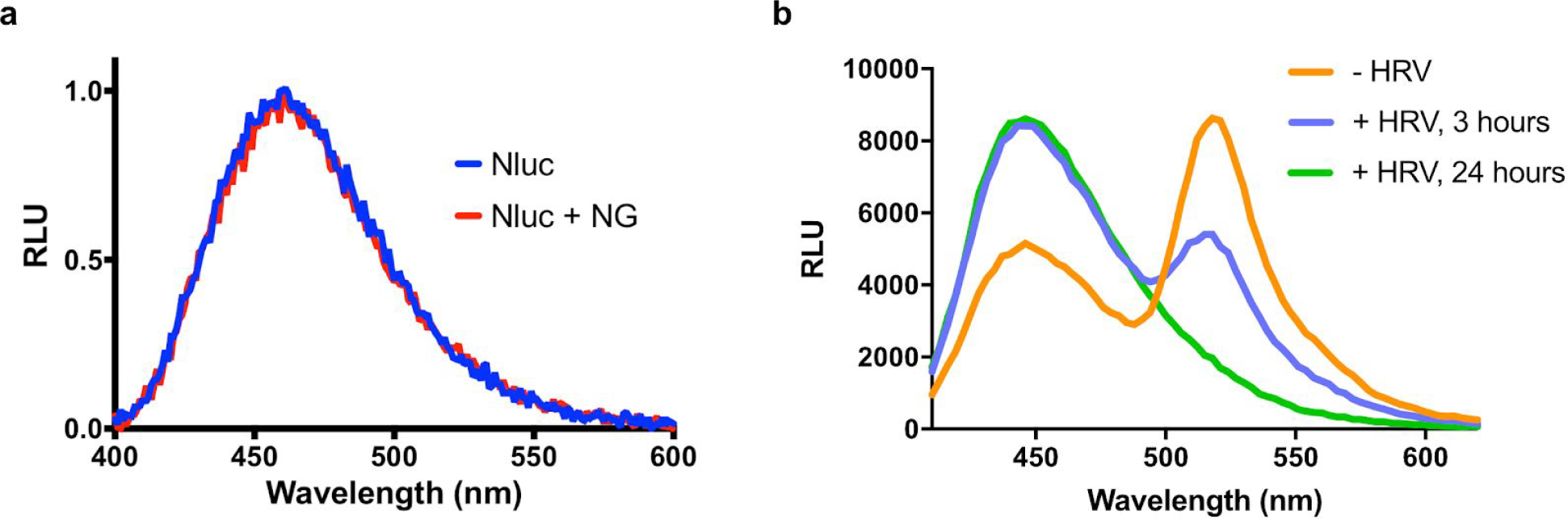
Validation of BRET tension sensor. (A) NanoLuc (Nluc) and mNeonGreen (NG) was inserted into a cell surface receptor. NanoLuc was transfected into mammalian cells either alone or in a cotransfection with mNeonGreen. Readout was performed on a plate reader. Units in relative light units (RLU). (B) A 3C protease site was cloned between NanoLuc and mNeonGreen. Recomibantly expressed protein was exposed to Human Rhinovirus (HRV) 3C protease and measured on a plate reader following the labeled time.

**Figure S3.**
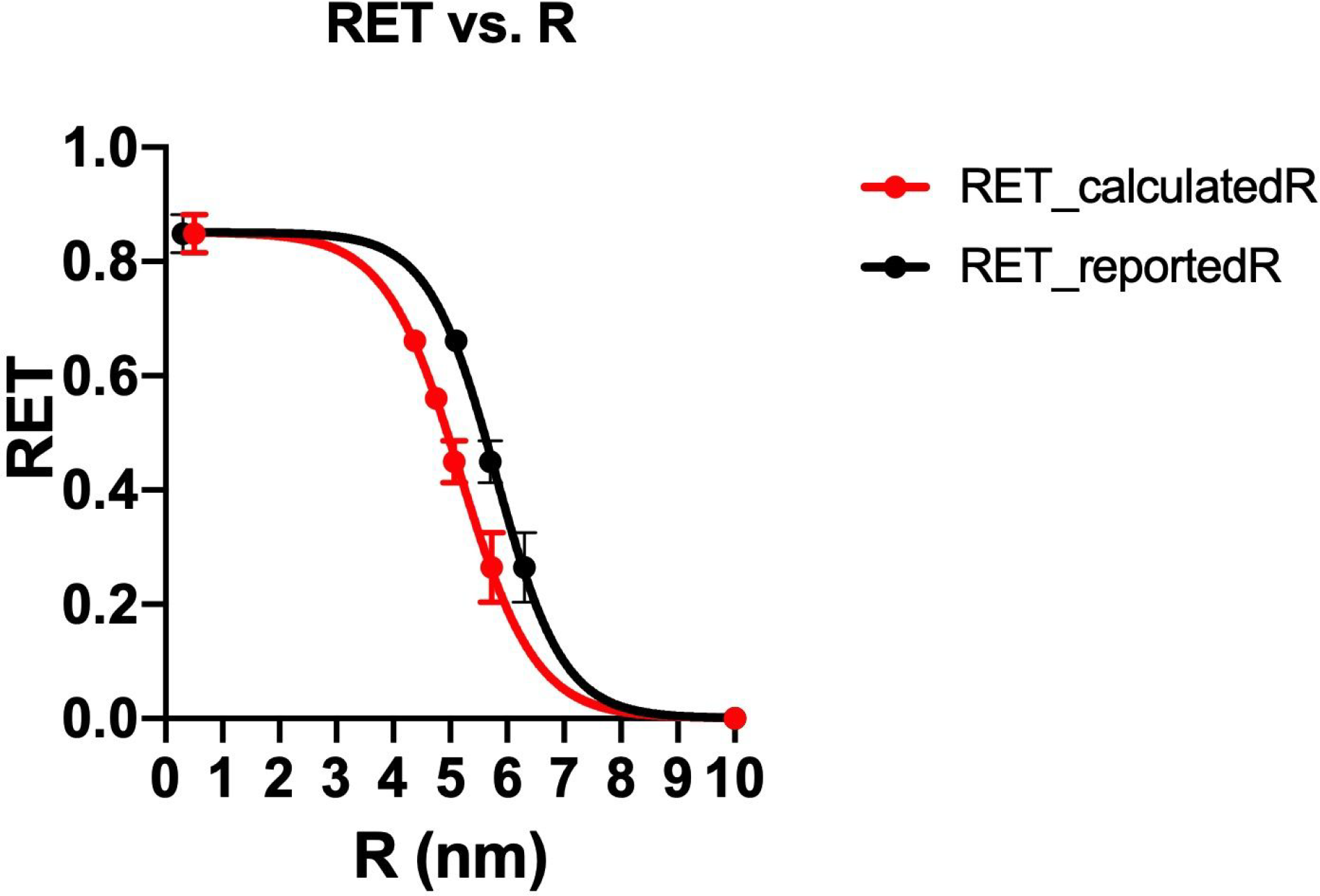
Resonance energy transfer (RET) efficiency curves resulting from R’s calculated in this work, using an R_0_ of 5.1nm, or R’s reported in the literature. Curve generated using calculated distance (R) values for the series of HL linkers separating mNeonGreen and NanoLuc or using reported HL linker distance values^19^. Error bars denote differences between acquiring data on recombinant protein or from mammalian cell surface expressed sensors.

**Figure S4.**
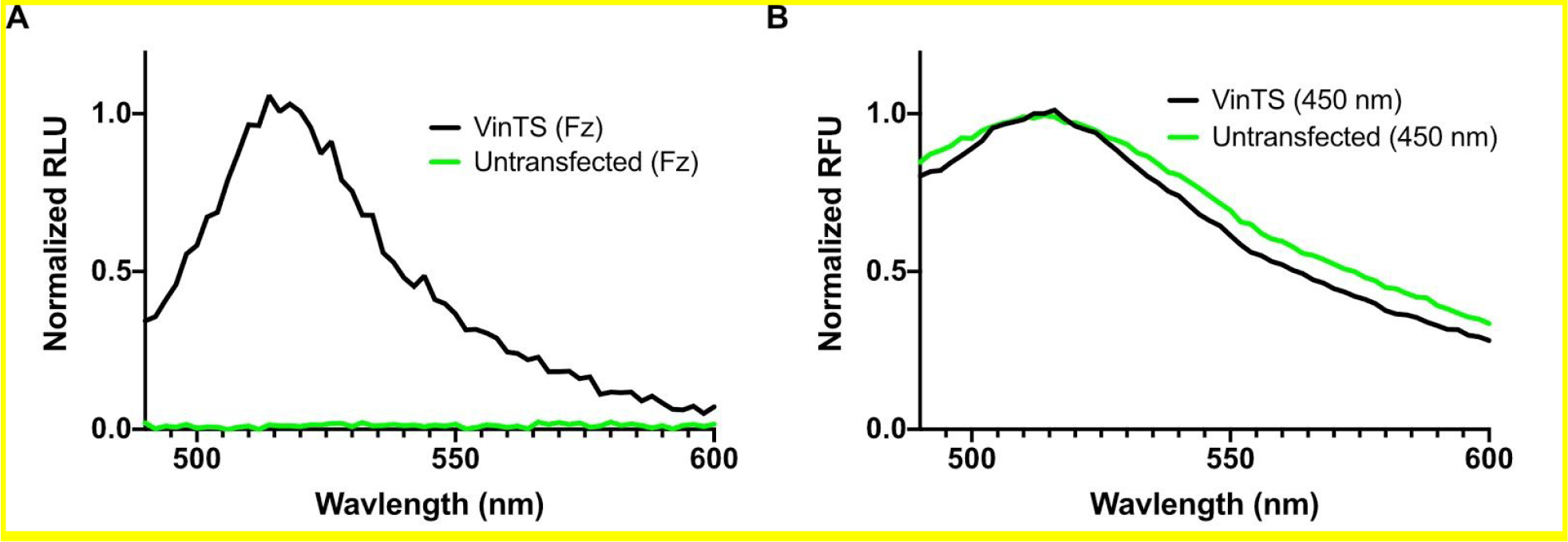
Comparison of mNeonGreen emission resulting from BRET or direct excitation. Emission of mNeonGreen of Untransfected or HEK293T cells transfected with VinTS was measured on a plate reader upon excitation with: (A) furimazine (Fz) to induce Luminescence. Units in relative luminescence units (RLU). Untransfected cells show no signal while VinTS expressing cells exhibit clear mNeonGreen emission (B) 450 nm light to induce fluorescence. Autofluorescence from cells exhibits same emission signal as excitation of mNeonGreen, underscoring the sensitivity and low background of BRET. Units in relative fluorescence units (RFU). Emissions normalized to 1. Experiment in triplicate (n=3).

**Figure S5.**
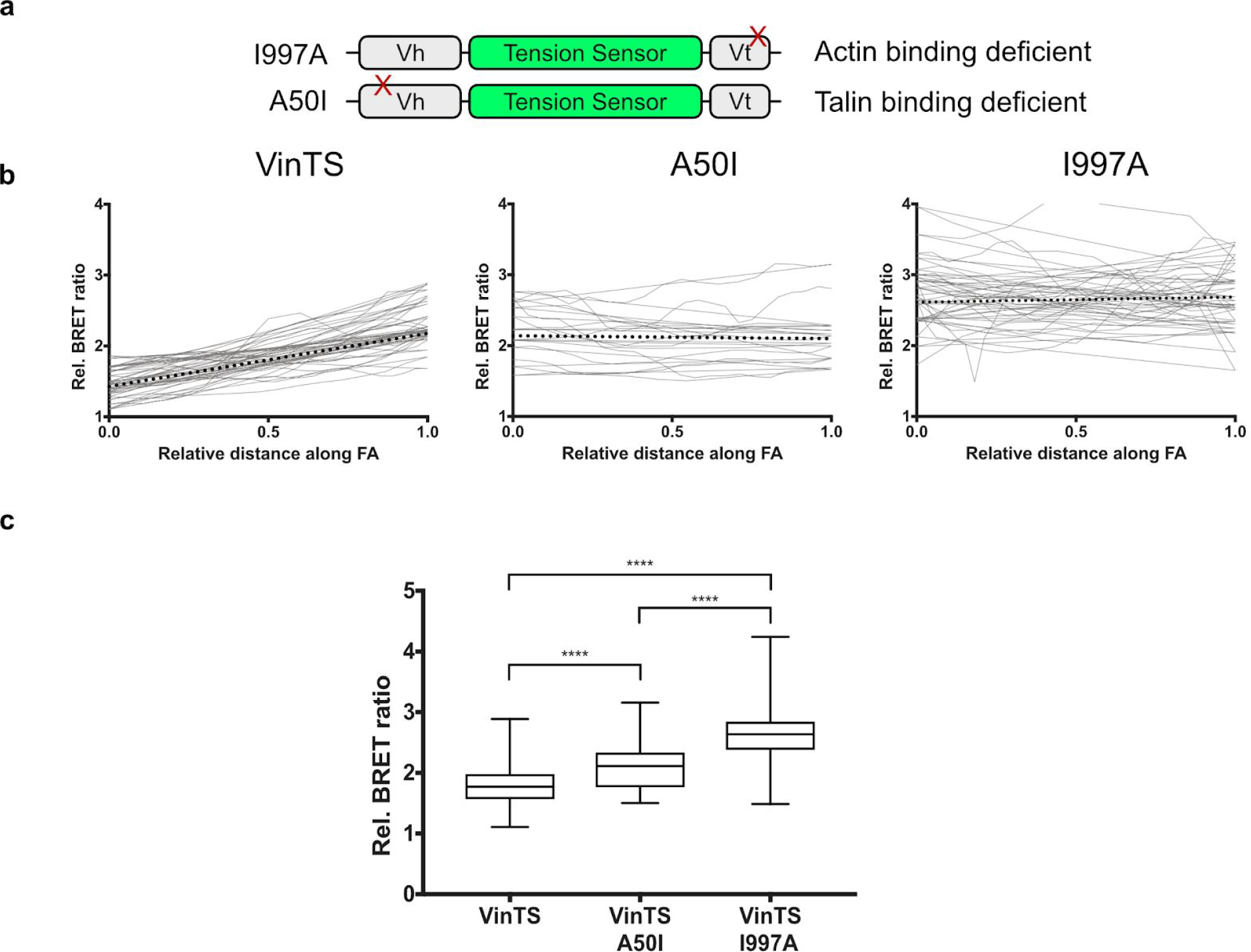
Point mutations in BRET-TS vinculin. (A) Schematic of point mutations made in vinculin to abolish either actin binding (I997A) or talin binding (A50I) with inserted BRET-TS^12^. (B) Representative line scans taken across focal adhesions (FA) in ratiometric images for the various constructs. N = 30 focal adhesions across 3 cells. Numerous independent experiments were performed and acquired similar data. (C) Quantified average relative BRET ratio from line scans in (B). Data analysis via 2-tailed Student’s t-test. Significance: **** p<0.001.

## Supplemental sequences

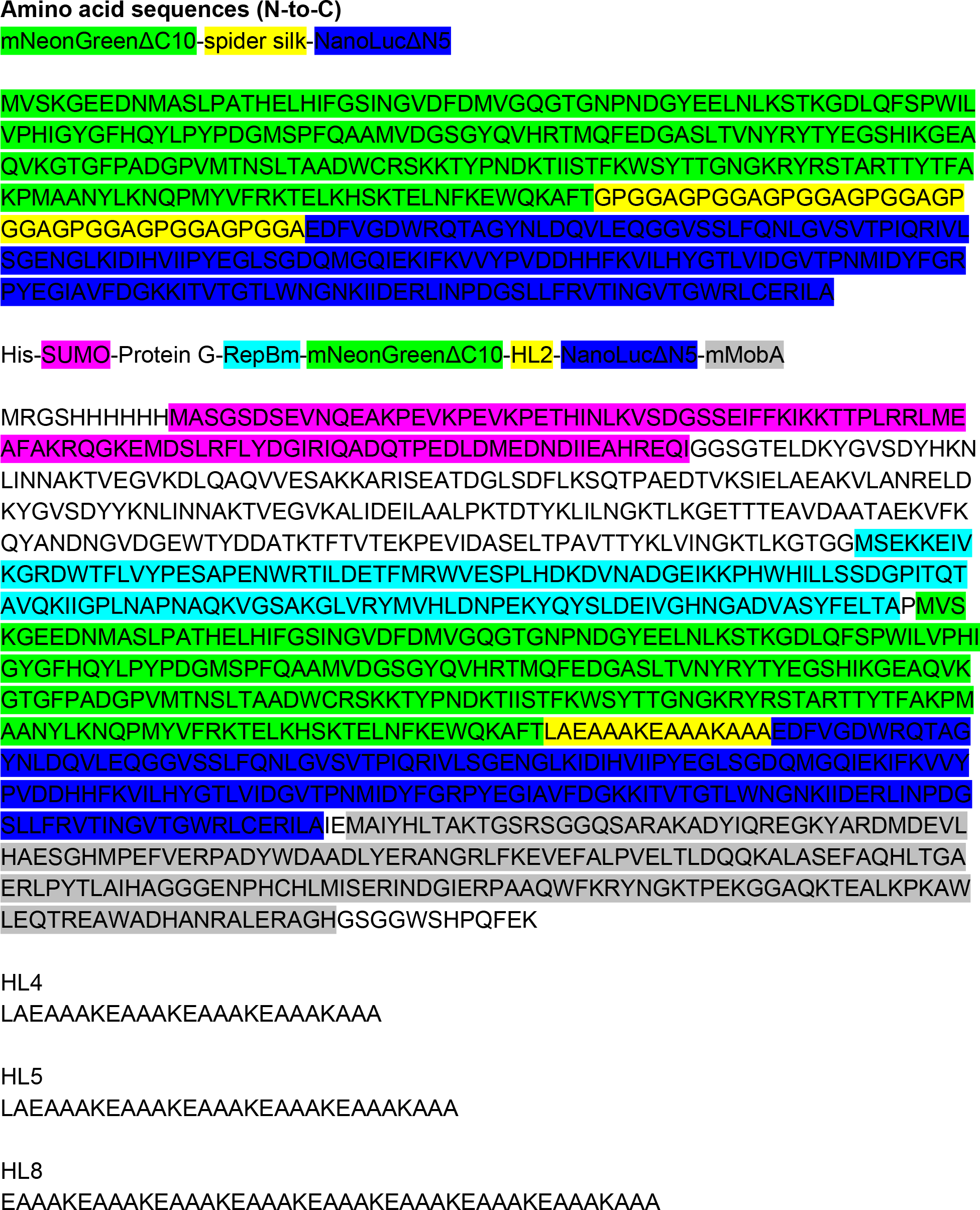

